# Genomic and phylogenetic analyses of SARS-CoV-2 strains isolated in the city of Gwangju, South Korea

**DOI:** 10.1101/2020.12.16.423178

**Authors:** Ji-eun Lee, Jae Keun Chung, Tae sun Kim, Jungwook Park, Mi hyeon Lim, Da jeong Hwang, Jin Jeong, Kwang gon Kim, Ji-eun Yoon, Hye young Kee, Jin jong Seo, Min Ji Kim

## Abstract

Since the first identification of severe acute respiratory syndrome coronavirus 2 (SARS-CoV-2) in China in late December 2019, the coronavirus disease 2019 (COVID-19) has spread fast around the world. RNA viruses, including SARS-CoV-2, have higher gene mutations than DNA viruses during virus replication. Variations in SARS-CoV-2 genome could contribute to efficiency of viral spread and severity of COVID-19. In this study, we analyzed the locations of genomic mutations to investigate the genetic diversity among isolates of SARS-CoV-2 in Gwangju. We detected non-synonymous and frameshift mutations in various parts of SARS-CoV-2 genome. The phylogenetic analysis for whole genome showed that SARS-CoV-2 genomes in Gwangju isolates are clustered within clade V and G. Our findings not only provide a glimpse into changes of prevalent virus clades in Gwangju, South Korea, but also support genomic surveillance of SARS-CoV-2 to aid in the development of efficient therapeutic antibodies and vaccines against COVID-19.

## Introduction

The coronavirus disease 2019 (COVID-19) is caused by a newly emerged virus, named severe acute respiratory syndrome coronavirus 2 (SARS-CoV-2) (1, 2). Since its first reporting on December 2019 in Hubei Province, China, COVID-19 has rapidly spread globally, and the World Health Organization (WHO) declared it a pandemic on March 11, 2020 (3). As of April 20, 2020, 2,995,758 human confirmed cases of COVID-19, including 204,987 deaths, have been reported from 213 countries (4). The first case in South Korea, a Chinese visitor from Wuhan, was identified on January 20, 2020 (5, 6). On February 3, 2020, the first case in the city of Gwangju, South Korea, was reported, and as of mid-July 2020, a total of 171 confirmed cases have been identified.

SARS-CoV-2 is an enveloped, positive-sense single-stranded RNA virus (7). The SARS-CoV-2 genome encodes 16 non-structural proteins (nsps) involved in virus replication and four structural proteins, including envelope (E), matrix (M) and nucleocapsid (N) and spike (S) (8). RNA viruses have higher mutation rates than DNA viruses (9). Mutations in the SARS-CoV-2 genome are being continuously reported (10, 11), and these mutations may affect pathogenesis. Therefore, it is critically important to monitor the genome evolution of SARS-CoV-2. Investigation of the viral genomic variations is necessary to provide epidemiological information on SARS-CoV-2 and for the development of therapeutics and vaccines.

In this study, we isolated SARS-CoV-2 virus from COVID-19 patients in the city of Gwangju, South Korea, and we investigated genomic mutations resulting in amino acid changes. To this end, we performed full-genome sequencing using a next-generation sequencing (NGS) tool and analyzed point mutations in the SARS-CoV-2 genome. We phylogenetically analyzed and classified virus strains from confirmed SARS-CoV-2-infected patients in the southwestern region of Korea.

## Materials and Methods

### Sampling and viral RNA isolation

This study was approved by the Institutional Review Board (approval no. P01-202008-31-004) of Public Institutional Bioethics Committee designated by the Ministry of Health of Welfare (MOHW). Clinical specimens were collected as oropharyngeal and nasopharyngeal swabs in viral transport medium or sputum from symptomatic patients with COVID-19. All clinical samples were handled in biosafety cabinets. Viral RNA was extracted from the samples using a QIAamp Viral RNA Mini Kit (Qiagen, Hilden, Germany). The RNA was quantified by the Korea Centers for Disease Control and Prevention (KCDC) method and using a PowerChek 2019-nCoV Real-time Kit (Kogene Biotech, Seoul, South Korea).

### Real-time reverse-transcription polymerase chain reaction (RT-qPCR)

RT-qPCR assays targeted the RNA-dependent RNA polymerase (*RdRp*) and *E* genes. Sequence information for the primers and probes used to detect these two genes is presented in Table 1 (12). PCRs were run in 25-μL reaction mixtures containing 12.5 μL of 2X RT-PCR buffer, 1 μL of 25X RT-PCR enzyme mixture included in AgPath-ID One-step RT-PCR Reagents (Thermo Fisher Scientific), 5 μL of RNA, and 1 μL of each primer/probe. The PowerChek 2019-nCoV Real-time PCR Kit includes a primer/probe mixture (for *RdRp* and *E*), a positive control template (for *RdRp* and *E*), and RT-qPCR premix. Sequences of the primers and probe are not disclosed. The RT-qPCR master mix consists of 11 μL of RT-qPCR premix, 4 μL of each primer/probe mixture, and 5 μL of RNA. The thermal cycling program was as follows: reverse transcription at 50°C for 30 min, followed by inactivation of the reverse transcription at 95°C for 10 min, and then 40 cycles of 95°C for 15 s and 60°C for 1 min (7500 instrument; Applied Biosystems, Foster City, CA, USA). All samples were tested in duplicate.

**Table 1.**
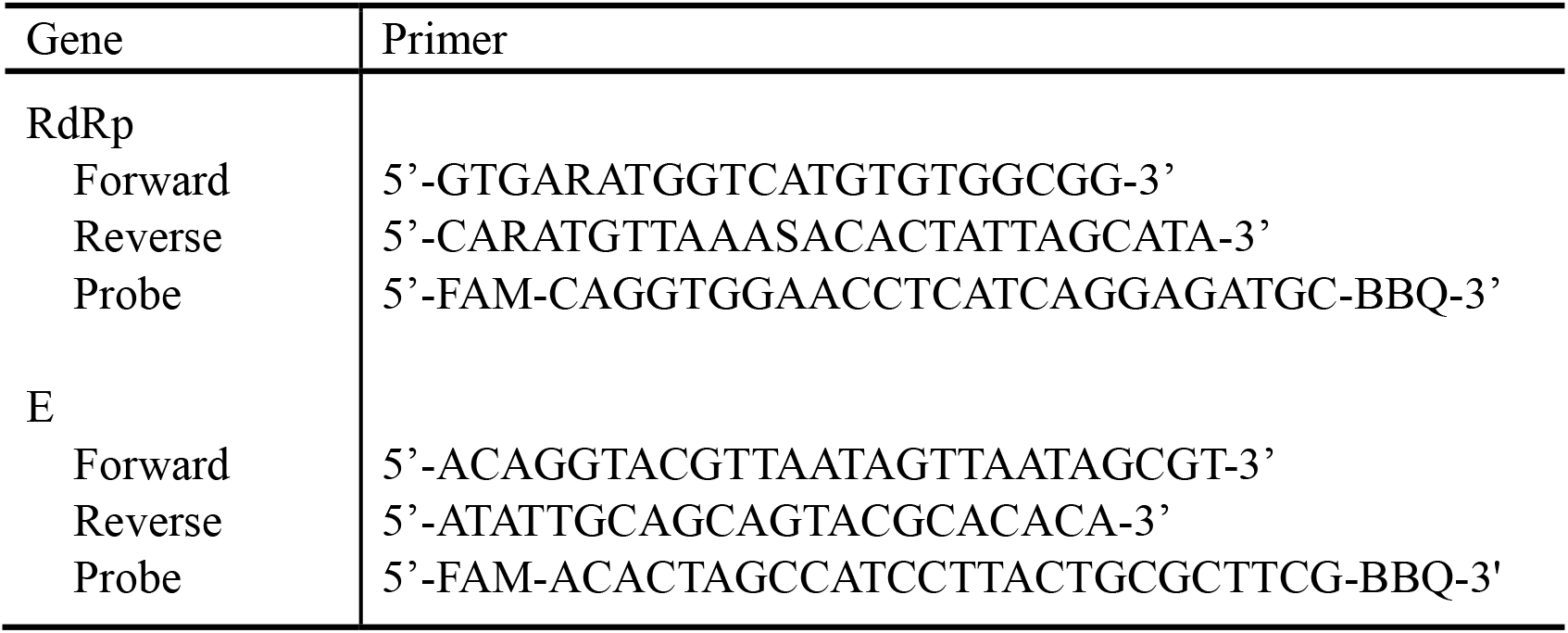
RT-qPCR primers/probes (KCDC)

### SARS-CoV-2 isolation from Vero cells

To isolate SARS-CoV-2 from Vero cells, the cells were seeded in 24-well plates at 2×10^4^ cells/well and cultured in modified Eagle’s medium (MEM; Gibco, Thermo Fisher Scientific, USA) supplemented with 10% heat-inactivated fetal bovine serum (FBS; Gibco) for 48 h. Then, the medium was replaced with medium with a lower FBS concentration, and the cells were further incubated for 24–48 h. Before inoculation onto the Vero cells, patient specimens were diluted with viral transport medium containing penicillin-streptomycin (Gibco) at a 1:4 ratio, incubated at 4°C 1 h with vortexing every 15 min, and then centrifuged at 3,000 rpm for 30 min. Inoculated monolayers of Vero cells were incubated in MEM supplemented with 2% FBS at 37°C in the presence of 5% CO_2_. For seven days post inoculation, the cells were observed for cytopathic effects (CPE) daily. When CPE were observed, the cells were scraped with a micropipette tip and subjected to total nucleic acid extraction.

### Next-generation sequencing

Ten cell supernatants containing viral particles and six clinical samples with a threshold cycle (Ct) value of 10 or lower were used for genome analysis. Viral RNA was isolated using the QIAamp Viral RNA Mini Kit and subjected to whole-genome sequencing. NGS libraries were prepared and sequenced using Barcode-Tagged Sequencing (BTSeq) technology (Celemics Inc., Seoul, Korea) according to the manufacturer’s protocol. Complete genome sequencing was conducted using the Illumina MiSeq platform (Illumina, San Diego, CA), producing 150-bp pair-end reads per sample.

### Phylogenetic analysis

Forty complete viral genome sequences were downloaded from the Global Initiative on Sharing All Influenza Data (GISAID; https://www.gisaid.org/) database, with acknowledgment. Multiple alignment of the SARS-CoV-2 genomes was carried out using Multiple Sequence Comparison by Log-Expectation (MUSCLE). A phylogenetic tree was generated by the neighbor-joining method using Molecular Evolutionary Genetics Analysis across Computing Platforms (MEGA X) using the Kimura 2-parameter model, and the tree evaluated using 1,000 bootstrap replicates.

### Mutation analysis

A reference SARS-CoV-2 genome (hCoV-19/Wuhan-Hu-1/2019) of 29,903 nucleotides long was obtained from the GISAID database and was used to identify mutations in the protein-coding sequences of the SARS-CoV-2 genomes of the Gwangju isolates. We analyzed 12 protein-coding sequences (ORF1a, ORF1b, S, ORF3a, E, M, ORF6, ORF7a, ORF7b, ORF8, N, and ORF10). The amino acid sequence was created by using hCoV-19/Wuhan-Hu-1/2019 annotated open reading frames (ORFs). Based on the protein annotations, amino acid codon variants were converted from the nucleotide variants for alignments. Each protein domain was aligned with the reference genome of SARS-CoV-2 using NCBI protein alignments.

## Results and Discussion

### Virus isolation from positive samples

For virus isolation, SARS-CoV-2-positive clinical specimens were inoculated into Vero cell cultures. We observed CPE at 24-h intervals for 7 days post inoculation. CPE were first observed and evident in Vero cells at 72–84 h after inoculation. When >80% of total cells showed CPE, the cells were collected (day 6), and viral RNA was extracted.

For the detection of SARS-CoV-2, the *RdRp* and *E* genes were detected using RT-qPCR. A positive RT-qPCR result was defined as a Ct value <37 for both genes. We successfully isolated SARS-CoV-2 from clinical samples collected from 10 COVID-19 patients in Gwangju. The virus isolates were named hCoV-19/Gwangju/IHC_sample number/2020 (Table 2).

**Table 2.**
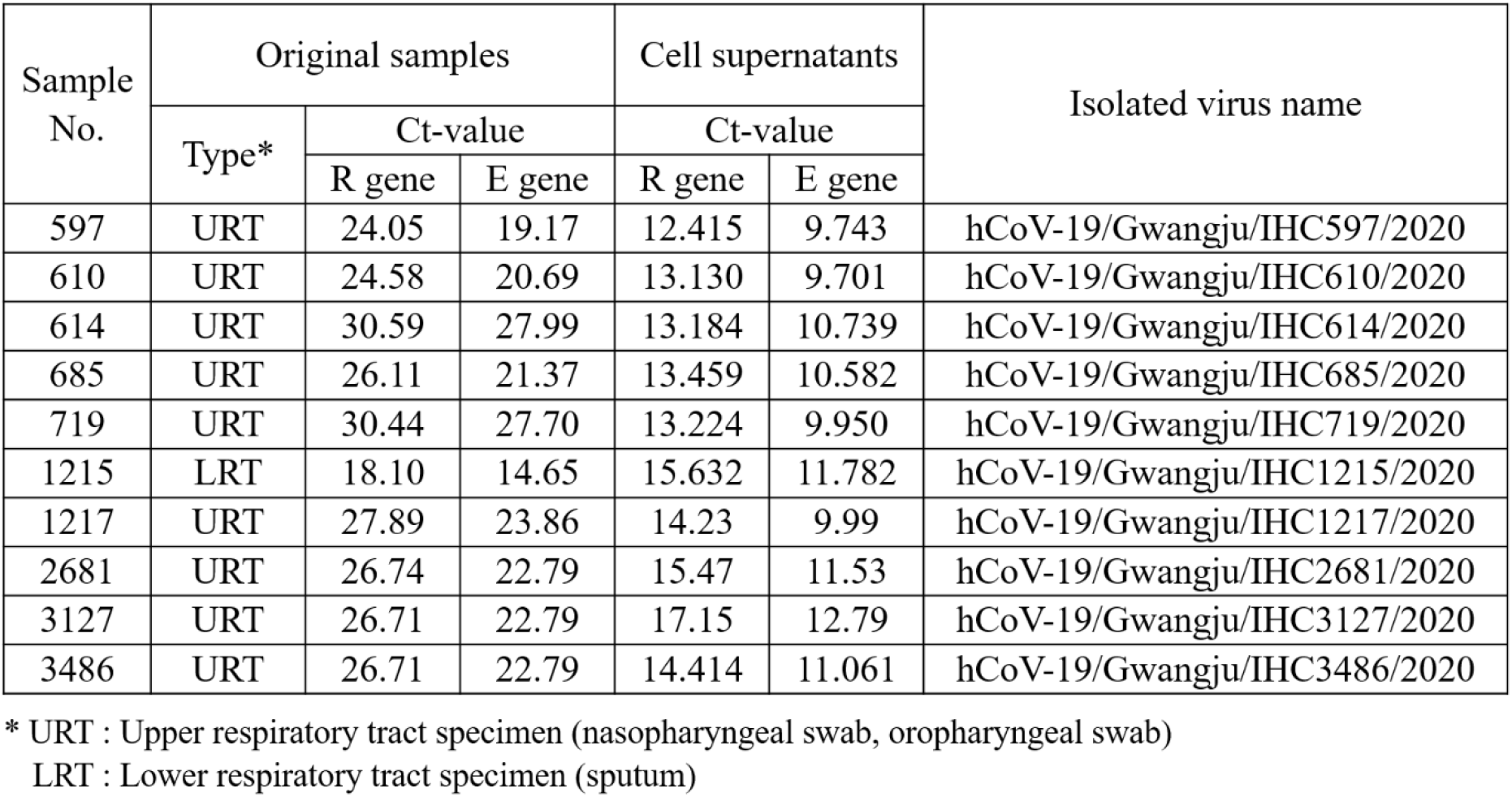
Isolated viral sample information

### Phylogenetic analysis of the SARS-CoV-2 genomes

We performed phylogenetic analyses of 16 distinct SARS-CoV-2 sequences of isolates from Gwangju and 40 sequences deposited in the GISAID database up to March 31, 2020 (Fig. 1). According to GISAID data, three major clades of SARS-CoV-2, including S, V, and G, can be identified (13). These three clades are determined by the amino acid substitutions present: ORF8-L84S (S), ORF3a-G251V (V), and S-D614G (G).

**Fig 1.**
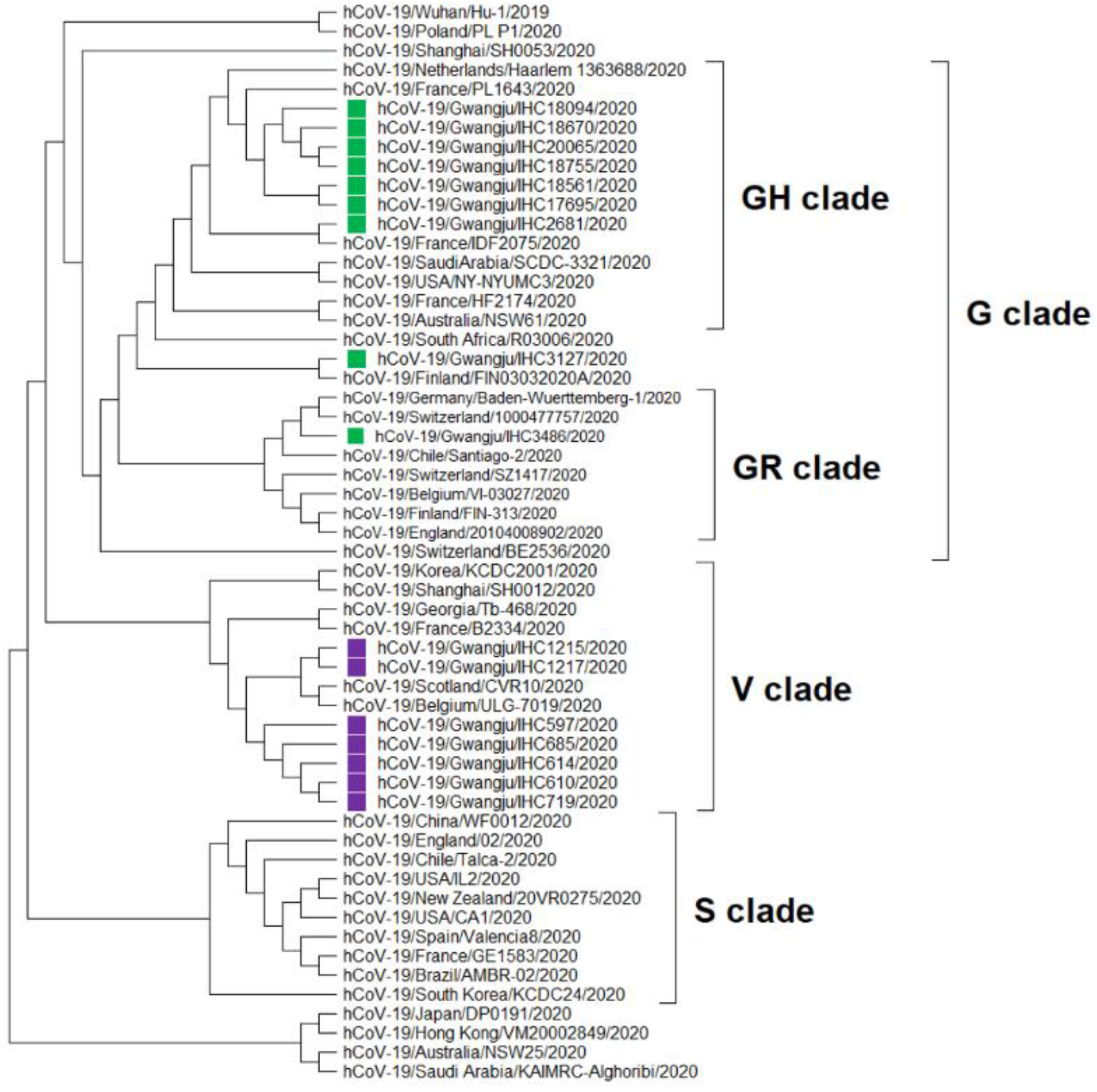
Phylogenetic analysis of 56 representative SARS-CoV-2 genomes, including sequences of viruses collected in the city of Gwangju, South Korea (green and purple). Clades S, V, and G were determined based on significant amino acid mutations (ORF8-L84S, ORF3a-G251V, and S-D614G, respectively). The G clade was classified into a GR clade, a GH clade, and other strains. Multiple sequence alignment was achieved using MUSCLE. The phylogenetic tree was constructed by the neighbor-joining method using MEGA X and the Kimura 2-parameter model. The best tree was found by performing 1,000 bootstrap replicates.

In accordance with the above classification, we identified three clades (GH, GR, and V) on the basis of the point mutations identified in the Gwangju sequences (Fig. 1). Clade V included seven isolates from Gwangju. Five of these isolates (indicated in purple in Fig. 1, IHC597, 610, 614, 685, and 719) were associated with the Shincheonji religious group, and the remaining two (Fig. 1, IHC1215, and 1217) were related to another church in Gwangju. As revealed by the epidemiologic information, the five isolates from this clade were related to the spread during the early COVID-19 pandemic in South Korea. Specifically, in February 2020, the COVID-19 outbreak started in the city of Daegu, South Korea, and SARS-CoV-2 had rapidly spread within the religious group named the Shincheonji church of Jesus (14). The G clade comprised nine of the 16 isolates (labeled green in Fig. 1). We identified two subclades within clade G: GH, with seven isolates, and GR, with one isolate. Of the seven confirmed cases related to the seven GH isolates, six were linked to cluster secondary infections from door-to-door sales businesses at Geumyang Building (Fig. 1, IHC17695, 18094, 18561, 18670, 18755 and 20065). A Geumyang Building case led to a cluster outbreak in Gwangju from late June to mid-July and further led to additional cases at the Gwangneuksa temple, church, and at work. The remaining one case appeared to be associated with international travel-related exposures (Fig. 1, IHC2681). The case related to the one isolate in clade GR had a known international travel history (Fig. 1, IHC3486). Taken together, these results reveal that the V clade of SARS-CoV-2 was the major clade type in Gwangju during the early pandemic in South Korea, and as of mid-July, clade GH is the predominant type in the southwestern region of South Korea.

### Mutations identified in the sequenced SARS-CoV-2 genomes

We aimed to classify the distinct clades of SARS-CoV-2 and investigate the mutations in the 16 SARS-CoV-2 genomes of the Gwangju isolates. To this end, we analyzed 12 protein-coding regions (ORF1a, ORF1b, S, ORF3a, E, M, ORF6, ORF7a, ORF7b, ORF8, N, and ORF10). NGS revealed 21 non-synonymous mutations and 1 frameshift deletion compared to the reference genome from a Wuhan isolate (NCBI GenBank: NC_045512). Mutation events observed in the sequenced SARS-CoV-2 genomes are summarized in Table 3.

**Table 3.**
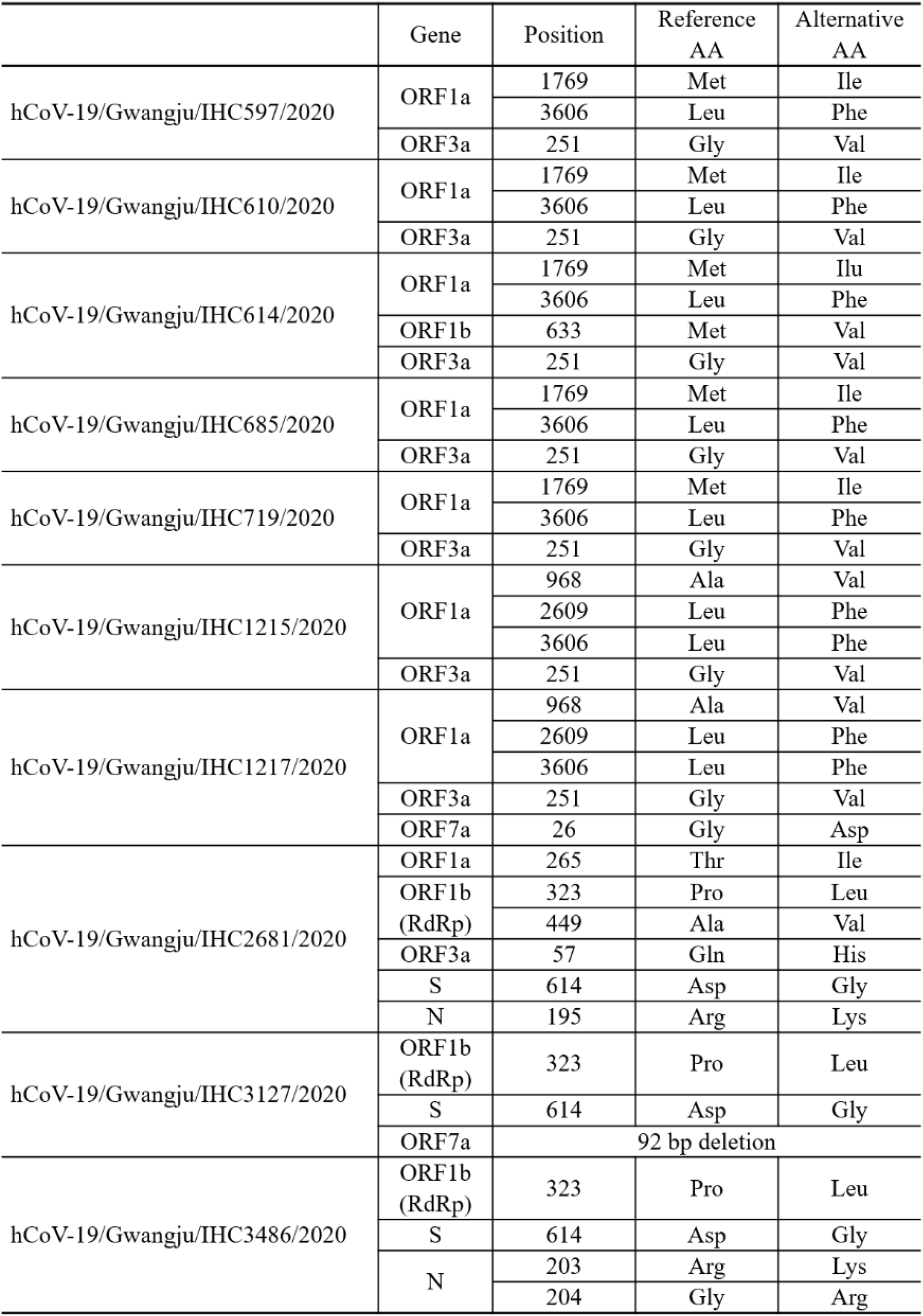
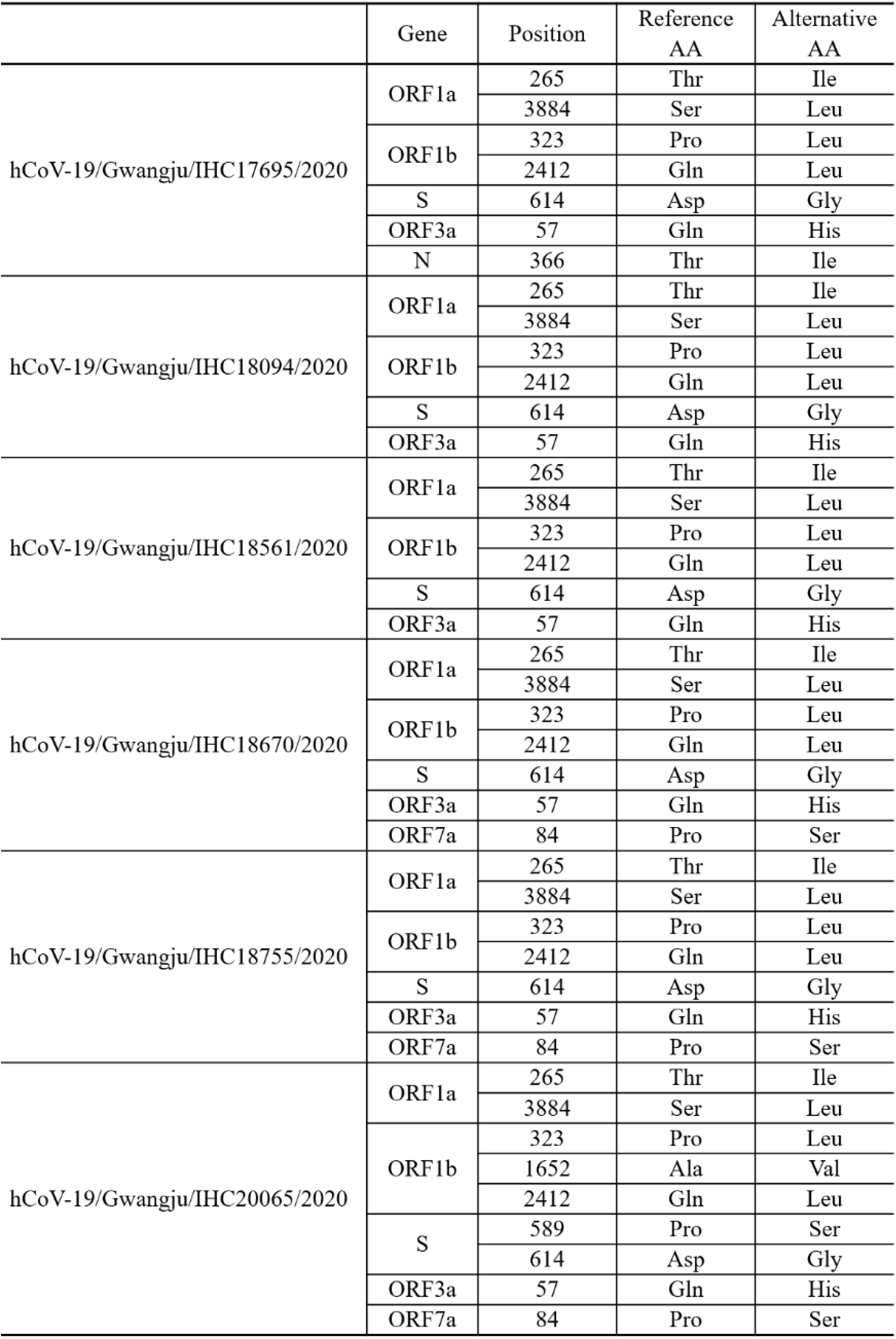
Amino acid mutations in the sequenced 16 SARS-CoV-2 genomes

Clades V (ORF3a-G251V) and G (S-D614G) comprised seven and nine out of the 16 sequences of Gwangju isolates, respectively (Table 4). ORF8-L84S, representative of clade S, was not observed in any of the 16 genomes. In addition to the G251V mutation in ORF3a and the D614G mutation in S, we analyzed 19 additional point mutations in ORF1a, ORF1b, ORF3a, ORF7a, S, and N. For these 19 genetic mutations, 11 mutation sites were found in ORF1a and ORF1b, which encode nsp1–16, occupying two-thirds of the entire genome (Fig. 2).

**Table 4.** Virus classification by representative mutations of each clade

| Virus name | Target gene<br>(reference AA) | ORF8<br>(L84) | ORF3a<br>(G251) | S<br>(D614) | Clade |
| --- | --- | --- | --- | --- | --- |
| hCoV-19/Gwangju/IHC597/2020 |  | L84 | V251 | D614 | V clade |
| hCoV-19/Gwangju/IHC610/2020 |  | L84 | V251 | D614 | V clade |
| hCoV-19/Gwangju/IHC614/2020 |  | L84 | V251 | D614 | V clade |
| hCoV-19/Gwangju/IHC685/2020 |  | L84 | V251 | D614 | V clade |
| hCoV-19/Gwangju/IHC719/2020 |  | L84 | V251 | D614 | V clade |
| hCoV-19/Gwangju/IHC1215/2020 |  | L84 | V251 | D614 | V clade |
| hCoV-19/Gwangju/IHC1217/2020 |  | L84 | V251 | D614 | V clade |
| hCoV-19/Gwangju/IHC2681/2020 |  | L84 | G251<br>(Q57H) | G614 | GH clade |
| hCoV-19/Gwangju/IHC3127/2020 |  | L84 | G251 | G614 | G clade |
| hCoV-19/Gwangju/IHC3486/2020 |  | L84 | G251 | G614 | GR clade |
| hCoV-19/Gwangju/IHC17695/2020 |  | L84 | G251<br>(Q57H) | G614 | GH clade |
| hCoV-19/Gwangju/IHC18094/2020 |  | L84 | G251<br>(Q57H) | G614 | GH clade |
| hCoV-19/Gwangju/IHC18561/2020 |  | L84 | G251<br>(Q57H) | G614 | GH clade |
| hCoV-19/Gwangju/IHC18670/2020 |  | L84 | G251<br>(Q57H) | G614 | GH clade |
| hCoV-19/Gwangju/IHC18755/2020 |  | L84 | G251<br>(Q57H) | G614 | GH clade |
| hCoV-19/Gwangju/IHC20065/2020 |  | L84 | G251<br>(Q57H) | G614 | GH clade |

**Fig 2.**
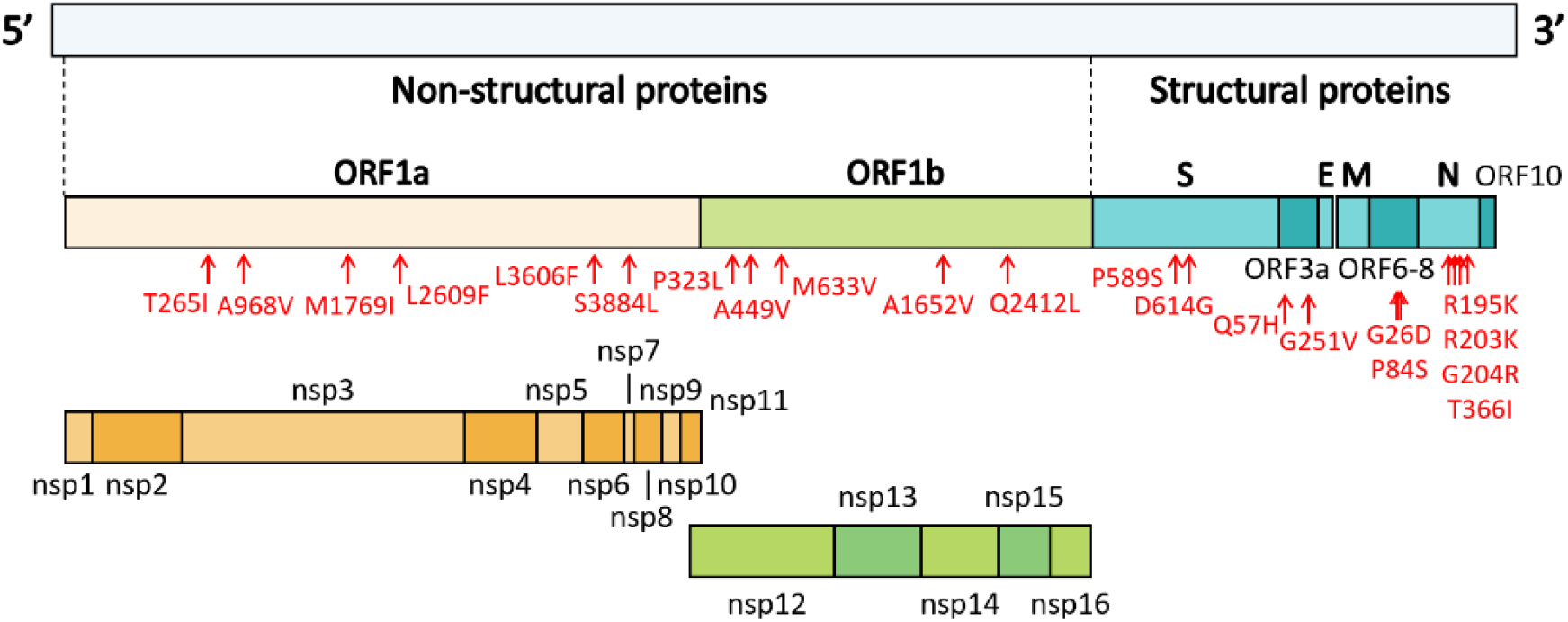
Genome organization of SARS-CoV-2. ORF1a and ORF1b encode 16 non-structural proteins (nsp1–16). The structural genes encode four structural proteins, including spike (S), envelope (E), matrix (M), and nucleocapsid (N). Mutation sites identified in the sequenced genomes of isolates from Gwangju are indicated in red.

We first analyzed the mutations in ORF1a/b, and we identified six mutation types in ORF1a (Table 3). Among them, the most common mutations were ORF1a-T265I (in seven samples, IHC2681, 17695, 18094, 18561, 18670, 18755, and 20065), ORF1a-L3606F (in seven samples, IHC597, 610, 614, 685, 719, 1215, and 1217) and ORF1a-S3884L (in six samples, IHC17695, 18094, 18561, 18670, 18755, and 20065). In particular, we found that ORF1a-L3606F co-occurred with ORF3a-G251V, which determines clade V (Table 3). Further, M1769I, A968V, and L2609F occurred in nsp3 within ORF1a. Mutations within nsp2 and nsp3 may play a role in differentiating infectivity of SARS-CoV-2 from SARS-CoV (15). In addition, we detected five mutations in ORF1b. P323L in nsp12 and Q2412L were observed in nine (IHC2681, 3127, 3486, 17695, 18094, 18561, 18670, 18755, and 20065) and six (IHC17695, 18094, 18561, 18670, 18755, and 20065) genome sequences, respectively. We also identified A449V and M633V in nsp12 and A1652V. The Nsp12 gene of SARS-CoV-2 encodes the RdRp protein essential for the virus replication machinery.

The remaining eight point mutations were found in ORF3a, ORF7a, S, and N. Four mutations, R195K, R203K, G204R, and T366I, were detected in the N protein, which participates in viral RNA genome packaging and viral particle release (16). Among them, G204R was observed together with ORF1b-P323L in the genome sequence of hCoV-19/Gwangju/IHC3486/2020 (Table 3). Based on GISAID data, two subclades, GR and GH, belonging clade G, were determined by two mutations (N-G204R and ORF1b-P323L) and one mutation (ORF3a-Q57H), respectively. This indicates that isolate hCoV-19/Gwangju/IHC3486/2020 can be classified into clade GR. Apart from the ORF3a-G251V and S-D614G substitutions, the single point mutations of Q57H in ORF3a and P589S in S were detected. In particular, ORF3a-Q57H, the signature of the GH clade, was detected in seven Gwangju genomes. In the ORF7a region, we confirmed two mutations; ORF7a-G26D and ORF7a-P84S.

In addition to the above 19 point mutations, we identified a 92-nucleotide deletion in ORF7a of hCoV-19/Gwangju/IHC3127/2020, resulting in out-of-frame deletion. The SARS-CoV ORF7a ortholog is an inhibitor of bone marrow stromal antigen 2 (BST-2), which is capable of restricting virus release from cells and induces apoptosis (17). According to this previous study, our observation implies that the deletion in ORF7a might affect the SARS-CoV-2 pathogenesis. However, further studies are needed to characterize the mechanisms underlying ORF7a functional alteration.

### Conclusions

We for the first time successfully isolated SARS-CoV-2 from COVID-19 patients in the southwestern region of South Korea. According to viral clade classification based on GISAID data, SARS-CoV-2 in Gwangju mainly belong to the V and G clades. More specifically, during the initial breakout in February 2020, the V clade was major clade type, mainly due to spread in the Shincheonji religious group. Since July 2020, the GH clade, related to door-to-door sales, became the prevalent type. In addition, we identified 21 non-synonymous mutations and one frameshift deletion. This study provided the first genomic mutation analysis in SARS-CoV-2 isolated from patients in Gwangju, South Korea. It is important to analyze and share the genome characteristics of SARS-CoV-2 for interpreting alterations in infectivity, pathogenicity, and host-virus interactions by continuous monitoring of the genetic mutations. Future studies including additional genome sequences are needed to define the further evolution of the SARS-CoV-2 epidemic in Gwangju.

## References

1. Pan A, Liu L, Wang C, Guo H, Hao X, Wang Q, Huang J, He N, Yu H, Lin X, Wei S, Wu T. 2020. Association of public health interventions with the epidemiology of the COVID-19 outbreak in Wuhan, China. JAMA 323:1915–1923.

2. Park SE. 2020. Epidemiology, virology, and clinical features of severe acute respiratory syndrome-coronavirus-2 (SARS-CoV-2; Coronavirus Disease-19). Clin Exp Pediatr 63:119–124.

3. World Health O. 2020. Coronavirus disease 2019 (COVID-19): situation report, 78. World Health Organization, Geneva.

4. Hui DS, I Azhar E, Madani TA, Ntoumi F, Kock R, Dar O, Ippolito G, McHugh TD, Memish ZA, Drosten C, Zumla A, Petersen E. 2020. The continuing 2019-nCoV epidemic threat of novel coronaviruses to global health — The latest 2019 novel coronavirus outbreak in Wuhan, China. Int J Infect Dis 91:264–266.

5. Kim JY, Choe PG, Oh Y, Oh KJ, Kim J, Park SJ, Park JH, Na HK, Oh M-d. 2020. The first case of 2019 novel coronavirus pneumonia imported into Korea from Wuhan, China: Implication for infection prevention and control measures. J Kor Med Sci 35:e61.

6. Kim J-M, Chung Y-S, Jo HJ, Lee N-J, Kim MS, Woo SH, Park S, Kim JW, Kim HM, Han M-G. 2020. Identification of coronavirus isolated from a patient in Korea with COVID-19. Osong Public Health Res Perspect 11:3–7.

7. Mousavizadeh L, Ghasemi S. 2020. Genotype and phenotype of COVID-19: Their roles in pathogenesis. J Microbiol Immunol Infect doi: https://doi.org/10.1016/j.jmii.2020.03.022.

8. Chan JF-W, Kok K-H, Zhu Z, Chu H, To KK-W, Yuan S, Yuen K-Y. 2020. Genomic characterization of the 2019 novel human-pathogenic coronavirus isolated from a patient with atypical pneumonia after visiting Wuhan. Emerg Microbes Infect 9:221–236.

9. Duffy S. 2018. Why are RNA virus mutation rates so damn high? PLOS Biology 16:e3000003.

10. Sheikh JA, Singh J, Singh H, Jamal S, Khubaib M, Kohli S, Dobrindt U, Rahman SA, Ehtesham NZ, Hasnain SE. 2020. Emerging genetic diversity among clinical isolates of SARS-CoV-2: Lessons for today. Infect Genet Evol 84:104330.

11. van Dorp L, Acman M, Richard D, Shaw LP, Ford CE, Ormond L, Owen CJ, Pang J, Tan CCS, Boshier FAT, Ortiz AT, Balloux F. 2020. Emergence of genomic diversity and recurrent mutations in SARS-CoV-2. Infect Genet Evol 83:104351.

12. Corman VM, Landt O, Kaiser M, Molenkamp R, Meijer A, Chu DK, Bleicker T, Brünink S, Schneider J, Schmidt ML, Mulders DG, Haagmans BL, van der Veer B, van den Brink S, Wijsman L, Goderski G, Romette J-L, Ellis J, Zambon M, Peiris M, Goossens H, Reusken C, Koopmans MP, Drosten C. 2020. Detection of 2019 novel coronavirus (2019-nCoV) by real-time RT-PCR. Euro surveillance 25:2000045.

13. Forster P, Forster L, Renfrew C, Forster M. 2020. Phylogenetic network analysis of SARS-CoV-2 genomes. Proc Natl Acad Sci U S A 117:9241–9243.

14. Park JY, Han MS, Park KU, Kim JY, Choi EH. 2020. First pediatric case of coronavirus disease 2019 in Korea. J Kor Med Sci 35:e124

15. Angeletti S, Benvenuto D, Bianchi M, Giovanetti M, Pascarella S, Ciccozzi M. 2020. COVID-2019: The role of the nsp2 and nsp3 in its pathogenesis. J Med Virol 92:584–588.

16. Narayanan K, Chen C-J, Maeda J, Makino S. 2003. Nucleocapsid-independent specific viral RNA packaging via viral envelope protein and viral RNA signal. J Virol 77:2922–2927.

17. Taylor JK, Coleman CM, Postel S, Sisk JM, Bernbaum JG, Venkataraman T, Sundberg EJ, Frieman MB. 2015. Severe acute respiratory syndrome coronavirus ORF7a inhibits bone marrow stromal antigen 2 virion tethering through a novel mechanism of glycosylation interference. J Virol 89:11820–11833.

